# Knowledge and Awareness of coronavirus disease 2019 (COVID-19) among Chinese dental students——a comparison study

**DOI:** 10.1101/2021.02.03.429522

**Authors:** Chengdan Deng, Huangshui Ma, Yuke Shou, Yuxuan Zhao, Yang Li, Bing Shi

## Abstract

**Backgroud:** This study aimed to measure the knowledge and awareness of COVID-19 among Chinese dental students during the global outbreak recently.

**Method:** A descriptive cross-sectional study was performed among dental students and nonmedical college students in China. All the participants were required to anonymously answer a reliable online questionnaire, which covered 3 different fields of COVID-19. Average scores of dental students (D group), including junior (JD group) and senior dental students (SD group), and nonmedical college students (N group) were compared respectively. Chi-square test and independent sample T test were taken for statistical analysis with SPSS.12.

**Results:** Totally 497 questionnaires were collected, including 224 from dental students and 273 from non-medical students. The overall average score was 57±19.2. The average scores of dental students were 64.5±18. The D group had significantly higher scores on the total score, section scores, and 20 questions respectively than with the N group. No significant differences were found on 5 questions. Compared with the N group, the SD group won on all three sections while JD group failed to win on the diagnose section.

**Conclusion:** Although the dental student showed good awareness regarding the clinical aspects of COVID-19 than non-medical students, there are still some weakness in the part of treatment and prevention, which need to be strengthened for better prepare during work. Besides, the low accuracy rate of lower grade dental students is also worth noting.

## Introduction

COVID-19 is a highly contagious viral infection that has caused global spread.[1, 2],More than 70,000 people have been confirmed infected in china [3], and the mortality is about 3.5%. Among those diagnosed, the ages range from 3 months to 91years old which shows people of all ages are susceptible. [4].And now that its spread to 6 continents and 52 countries and since February 25th the number of confirmed cases has risen rapidly in South Korea, Italy. [5][6] Some draconian public-facing measures have been taken in these countries.

More notable is the possibility of cross-infection in hospitals. From January 1 to 28, 29% were medical workers among the 138 patients admitted to Zhong Nan hospital of Wuhan university[7]. Faced with the newly discovered infectious disease, health workers are at high risk of infection. At the same time, as the first contact with the COVID-19, onset time of affected health workers is earlier than the social population, which forms one of the transmission chain of the outbreak. The currently available evidence on the modes of transmission is determined transmission by droplets and close contact with infected individuals or surface of equipment. As also, health workers should take greater precautions in aerosol-generating procedures.

The dentists are facing a high risk for COVID-19 infection due to the close contact with patients and facing the patient’s nose and mouth making the spray of droplets (patient’s blood and saliva) are difficult to avoid during treatment. Because the use of ultrasonic scaling in oral cavity and high-speed turbine phones can generate aerosols, which can increase the risk of virus transmission. Therefore, Senior dental students have direct contact with all patients during the internship. Do they have enough knowledge to protect them from infection? We designed an online questionnaire to find out how well oral medical students across the country knew about covid-19.

## Methods

### 1. Preliminary study

This descriptive cross-sectional study used convenience sample. Fifty dental students and fifty non-medical students were invited to participate in the preliminary study. The average accuracy is 67.54% in dental students and 54.34% in non-medical students. With a margin of error of 5%, power of study of 80%, 95% confidence level in the results, the sample size required at least 185 individuals for each group. According to the preliminary study, the Cronbach’s alpha was 0.7, which showed a well reliability of this questionnaire. The presentation and validity of the questionnaire were undertaken by experienced stomatology teacher and senior dental students.

### 2. Data collection

Data were collected between 19th −24th February 2020 at university students coming from 16 provinces in China. A total of 504 questionnaires were received, of which 497 were valid, and the effective rate was 98.6%,224 students in stomatology and 273 non-medical students as the control group. Their demographic characteristics were shown in Table1.

**Table 1.**
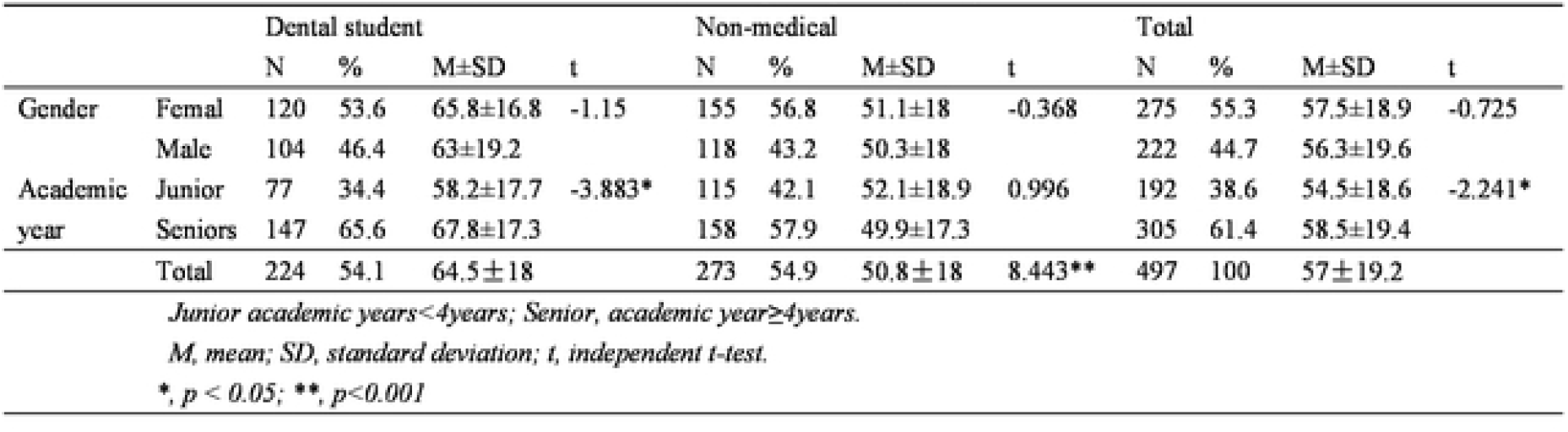
Baseline characteristics of the participants (n = 497).

According to the diagnosis and treatment protocols for COVID-19(6th edition) by National Health Commission of the People’s Republic of China, 20 true and false questions were designed as questionnaires and distributed online 【Table2】. These questions involved etiology, symptoms, and treatment, measurements for infection control and protection of COVID-19. All responses were anonymous and the basic information including gender, level of education, occupation and college will not be made public, so no ethics application has been made.

**Table 2.** Knowledge related to COVID-19(n= 497)

| Table2.Knowledge related to COVID-19(n=497) |  |  |  |  |  |  |  |
| --- | --- | --- | --- | --- | --- | --- | --- |
| Question(True or False) |  | A | D | N | J | S | Total |
| Transmission |  |  | N (%) | N (%) | N (%) | N (%) | N (%) |
| 1 | The digestive tract has been identified as a transmission | F | 188(83.9) | 243(89) | 65(84.4) | 123(83.7) | 431(86.7) |
| 2 | Transmission is mainly by respiratory tract and contact | T | 174(77.7) | 212(77.7) | 61(79.2) | 113(76.9) | 386(77.7) |
| 3 | The novel coronavirus nucleic acid could be detected in the fecal samples | T | 183(81.7) | 169(61.9) | 59(76.6) | 124(84.4) | 352(70.8) |
| 4 | Transportation by closed air-conditioning train belongs to close contact | T | 185(82.6) | 205(75.1) | 67(87) | 118(80.3) | 390(78.5) |
| 5 | Taking a Flight belongs to close contact | T | 109(48.7) | 94(34.4) | 36(46.8) | 73(49.7) | 203(40.8) |
| 6 | Transportation by airy bus belongs to close contact | F | 132(58.9) | 129(47.9) | 42(54.5) | 90(61.2) | 261(52.5) |
| Prevention |  |  |  |  |  |  |  |
| 7 | Chlorhexidine can effectively inactivate the fire virus | F | 162(72.3) | 150(54.9) | 48(62.3) | 114(77.6) | 312(62.8) |
| 8 | Chlorine-containing disinfectants can inactivate the virus | T | 191(85.3) | 199(72.9) | 61(79.2) | 130(88.4) | 390(78.5) |
| 9 | Lopinavir/ritonavir are identified as antiviral methods | F | 156(69.6) | 134(49.1) | 43(55.8) | 113(76.9) | 290(58.4) |
| 10 | Once the temperature returns to normal, the isolation can be released | F | 34(15.2) | 21(7.7) | 8(10.4) | 26(17.7) | 55(11.1) |
| 11 | Operations that may produce acrosols for diagnosed patients should be air-isolated | T | 172(76.8) | 149(54.6) | 56(72.7) | 116(78.9) | 321(64.6) |
| 12 | Operations for diagnosed patients should be performed in a closed room | F | 151(67.4) | 120(44) | 48(62.3) | 103(70.1) | 271(54.5) |
| 13 | Wash your hands in a basin for 30 seconds before eating. | F | 147(65.6) | 138(50.5) | 48(62.3) | 99(67.3) | 285(57.3) |
| 14 | Knives for raw and cooked food should be kept separate | T | 167(74.6) | 183(67) | 57(74) | 110(74.8) | 350(70.4) |
| Diagnosis |  |  |  |  |  |  |  |
| 15 | Mild cases have low fever, mild fatigue but no pneumonia | T | 177(79) | 171(62.6) | 50(64.9) | 127(86.4) | 348(70) |
| 16 | The mild cases had no symptoms of nasal congestion, runny nose, sore throat and diarrhea | T | 139(62.1) | 106(38.8) | 34(44.2) | 105(71.4) | 245(49.3) |
| 17 | Bilateral multiple lobula were observed on chest CT in early stage | F | 152(67.9) | 125(45.8) | 42(54.5) | 110(74.8) | 277(55.7) |
| 18 | Changes in intra-pulmonary would be found on chest CT in early stage | F | 94(42) | 53(19.4) | 25(32.5) | 69(46.9) | 147(29.6) |
| 19 | Lung becomes ground glass, infiltrating shadow early | F | 150(67) | 132(48.4) | 46(59.7) | 104(70.7) | 282(56.7) |
| 20 | Bilateral subsegmental areas of consolidation occurs early | F | 122(54.5) | 92(33.7) | 31(40.3) | 91(61.9) | 214(43.1) |
| 21 | A history of wuhan exposure is required for diagnosis | F | 73(32.6) | 69(25.3) | 18(23.4) | 55(37.4) | 142(28.6) |
| 22 | The diagnose in wuhan are different from others | T | 103(46) | 104(38.1) | 33(42.9) | 70(47.6) | 207(41.6) |
| 23 | The mild case has fever, cough and pneumonia | F | 161(71.9) | 169(61.9) | 51(66.2) | 110(74.8) | 330(66.4) |
| 24 | The normal case has fever and pneumonia | T | 128(57.1) | 131(48) | 41(53.2) | 87(59.2) | 259(52.1) |
| 25 | Severe cases may not have significant fever | T | 161(71.9) | 169(61.9) | 51(66.2) | 110(74.8) | 330(66.4) |
| A, answer,D, dental student; N, non-medical student; J, junior of dental student; S, senior of dental student. |  |  |  |  |  |  |  |
| N frequency, % percentage |  |  |  |  |  |  |  |

### 3. Statistical analysis

A correct answer was assigned 1 point and an incorrect answer were assigned 0 points. The total score was converted into a percentile. According to general characteristics, independent t-test was used to analysis difference of average score 【Table1】 and difference of three sections between two groups.【Table4】

**Table 3.** Comparison of the accuracy.

| Q | D/N |  | J/S |  | N/J |  | N/S |  |
| --- | --- | --- | --- | --- | --- | --- | --- | --- |
|  | × 2 | p | × 2 | p | × 2 | p | × 2 | p |
| 1 | 2.76 | 0.097 | 0.021 | 0.886 | 1.201 | 0.273 | 2.43 | 0.119 |
| 2 | 0 | 0.995 | 0.161 | 0.688 | 0.086 | 0.77 | 0.034 | 0.854 |
| 3 | 23.325 | <0.001** | 2.019 | 0.155 | 5.73 | 0.017* | 22.827 | <0.001** |
| 4 | 4.094 | 0.043* | 1.597 | 0.206 | 4.928 | 0.026* | 1.444 | 0.23 |
| 5 | 10.31 | 0.001* | 0.171 | 0.679 | 3.905 | 0.048* | 9.25 | 0.002* |
| 6 | 6.727 | 0.009* | 0.931 | 0.335 | 1.278 | 0.258 | 7.475 | 0.006* |
| 7 | 15.899 | <0.001** | 5.843 | 0.016* | 1.336 | 0.248 | 20.914 | <0.001** |
| 8 | 11.152 | 0.001* | 3.416 | 0.065 | 1.259 | 0.262 | 13.598 | <0.001** |
| 9 | 21.399 | <0.001** | 10.567 | 0.001* | 1.098 | 0.295 | 30.455 | <0.001** |
| 10 | 7.007 | 0.008* | 2.09 | 0.148 | 0.575 | 0.448 | 9.604 | 0.002* |
| 11 | 26.529 | <0.001** | 1.084 | 0.298 | 8.152 | 0.004* | 24.296 | <0.001** |
| 12 | 27.299 | <0.001** | 1.375 | 0.241 | 8.13 | 0.004* | 26.16 | <0.001** |
| 13 | 11.432 | 0.001* | 0.562 | 0.453 | 3.352 | 0.067 | 10.965 | 0.001* |
| 14 | 30341 | 0.068 | 0.017 | 0.896 | 1.363 | 0.243 | 2.754 | 0.097 |
| 15 | 15.727 | <0.001** | 14.035 | <0.001** | 0.136 | 0.712 | 26.166 | <0.001** |
| 16 | 26.555 | <0.001** | 15.962 | <0.001** | 0.71 | 0.399 | 40.621 | <0.001** |
| 17 | 24.291 | <0.001** | 9.533 | 0.002* | 1.847 | 0.174 | 32.7 | <0.001** |
| 18 | 30.039 | <0.001** | 4.345 | 0.037* | 5.909 | 0.015* | 35.124 | <0.001** |
| 19 | 17.366 | <0.001** | 2.768 | 0.096 | 3.117 | 0.077 | 19.47 | <0.001** |
| 20 | 21.638 | <0.001** | 9.546 | 0.002* | 1.134 | 0.287 | 30.916 | <0.001** |
| 21 | 3.226 | 0.072 | 4.533 | 0.033* | 0.116 | 0.734 | 6.768 | 0.009* |
| 22 | 3.149 | 0.076 | 0.461 | 0.497 | 0.572 | 0.45 | 3.572 | 0.059 |
| 23 | 5.482 | 0.019* | 1.847 | 0.174 | 0.482 | 0.487 | 7.158 | 0.007* |
| 24 | 4.135 | 0.042* | 0.727 | 0.394 | 0.665 | 0.415 | 4.8 | 0.028* |
| 25 | 5.482 | 0.019* | 1.847 | 0.174 | 0.482 | 0.487 | 7.158 | 0.007* |

*Q*, question; *D*, dental student; *N*, non-medical student; *J*, junior of dental student; *S*, senior of dental student.
$\chi^2$ , Chi-square, *p*, *p*-value,
\*: significance at < 0.05; \*\*: significance at < 0.001

**Table 4.** The score of different sections.

| Table4.The score of different sections. |  |  |  |  |  |  |  |  |
| --- | --- | --- | --- | --- | --- | --- | --- | --- |
|  | D/N |  | J/S |  | N/J |  | N/S |  |
|  | t | p | t | p | t | p | t | p |
| Transmission | 3.833 | <0.001** | -0.689 | 0.492 | -2.227 | 0.027* | -3.68 | <0.001** |
| Prevention | 6.255 | <0.001** | -2.175 | 0.031* | -2.694 | 0.007* | -6.862 | <0.001** |
| Diagnosis | 6.353 | <0.001** | -4.574 | <0.001** | -1.345 | 0.179 | -7.929 | <0.001** |
| *D, dental student; N, non-medical student; J, junior of dental student; S, senior of dental student.* |  |  |  |  |  |  |  |  |
| *t, independent t-test; p, p-value,* |  |  |  |  |  |  |  |  |
| *\*: significance at < 0.05; \*\* significance at < 0.001* |  |  |  |  |  |  |  |  |

The number and percentages were used to describe different categorical variables. 【Table2】The chi-square test was performed to examine the difference in accuracy between two groups (dental students and non-medical students). A reported p-value <0.05 was used to indicate statistical significance. 【Table3】

## Results

The dental students were significantly different with non-medical students on the total score and 20 questions. However, there was no difference between them on 5 questions.

Compared with junior in dental students, senior dental students had a higher accuracy on 18 questions and non-medical students had a lower accuracy on 6 questions. The senior in dental students had a higher accuracy than non-medical students on 17 questions (p-value <0.05).

On the compare of different sections, dental students were better than non-medical students in transmission, prevent ion and diagnosis, so were senior students and non-medical students. But junior students showed the same lack of knowledge as non-medical students on diagnosis of COVID-19

## Discussion

Although the dentists has many occupational exposure factors[8], their mortality rate is lower than normal, which may be due to the HWE[9, 10].In addition to the strategy of non-emergency delay, a better protective measures because of the higher awareness of the disease may also account for lower infection rates among health care workers[11]. In the knowledge of MERS investigation, the scores of medical students were significantly higher than that of nurses and other medical specialties which was same with this study. [12, 13]Previous medical background and knowledge could help a better understanding of COVID-19. In this study, although dental students showed a better understanding of COIVD-19, there were still some weaknesses that will be discussed below.

### 1. Transmission

More than 80% knew that droplets and close contact were the primary means of transmission. Interestingly, dental students were less correct on fecal-oral transmission than non-medical students, and so were seniors lower than junior. A latest report claimed that the SARS-Cov-2 was isolated from the feces of patients. But, there are still no case could confirm the fecal-oral transmission. Even so, the wrong answer may indicate that senior dental student pay a close attention to the updating news of COVID-19.

As a particular transmission during the oral clinic, 5% dental students admitted they didn’t know about aerosols. Moreover,17% didn’t know the correct way to avoid aerosol production. According to previous researches, MERS was easily spread by aerosols in medical facilities with central air conditioning, and had led to super-outbreaks of hospital-acquired infections[14].Prolonged exposure to high concentrations of aerosols in a closed environment may contribute to the spread of novel coronavirus[15]. The high speed turbine and ultrasonic tooth cleaning machine in the oral treatment tools will produce a large number of bioaerosols mixed with patients’ saliva and blood in the contaminated area from the patient’s head to the radius of the doctor’s back [16].Droplets mix in the air to form aerosols that can cause infection when inhaled[16]. There is no single way to reduce aerosol infections for dentist. It must be protected through personal protection, the use of rubber barrier, high efficiency indoor air filter, strong suction saliva, ultraviolet disinfection ventilation system and preventive mouthwash before treatment.[17-19] Thus, dental students need to strengthen understanding of aerosols to do a better protection.

To prevent close contact is in addition to the diagnosis and treatment process, more closely related to daily life.30%-50.5% dental senior students who had the highest accuracy still don’t unable to correctly determine which is close contact, which is worthy of caution. In the use of vehicles, people should try to avoid closed space such as fully closed air conditioning trains, buses, etc. Instead, well-ventilated and low-density vehicles should be used.

Hand hygiene is one of measures WHO recommendation for protection[2]. 35% dental student didn’t know it’s running water that wash hands with, not water in a sink, although most of them know when to wash their hands (before meals, after coughing, defecating, and touching animals).

### 2. Prevention

Choosing the appropriate disinfectant to inactivate the virus is necessary for every dentist. Research showed that dental students had a poor understanding of the control of infectious diseases,and so was this study. Coronavirus was sensitive to uv rays and could be effectively inactivated by 75% ethanol, chlorine-containing disinfectants or exposed for 30 minutes at 56 °C environment.23% dental students thought chlorhexidine could deactivate the coronavirus, which was wrong. 83% were not aware of the criteria for contact isolation which were normal temperature for more than 3 days, respiratory symptoms significantly improved better, inflammatory absorption in lung and negative detection of pathogenic nucleic acid.

### 3. Diagnosis

There is new evidence that afebrile but confirmed biologically patients can also spread the virus[20],which makes it more difficult for oral workers to identify infected patients as they work., it is also important for dentists to master the diagnostic conditions of COVID-19 to have a high level of clinical suspicion.

Mild cases with low fever, mild fatigue and no pneumonia are contagious and also the most likely to encounter in oral clinic. However, 40%dental student didn’t know what mild symptoms are, which is worth to watch out. The normal cases have fever, respiratory symptoms, and CT shows pneumonia. Although chest imaging was identified in the sixth edition of the protocol, nearly half of the dental seniors didn’t know it, this may due to the infrequent use of chest radiographs in dental clinics.

## Conclusion

Taking the right measures can better prevent and control the spread of COVID-19.Dental seniors generally master more knowledge of COVID-19 than ordinary students, there are still some deficiencies in relatively important treatment and prevention, which need to be strengthened in order to better prepare for work. In addition to medical students, ordinary college students, as a group receiving higher education, should also strengthen their knowledge about the epidemic and protect themselves and their families, so as to control the spread of the disease as soon as possible.

## Funding

The author(s) received no specific funding for this work.;

## Conflicts of interest

The authors declare that there is no conflict of interests regarding the publication of this paper.

